# Genome analysis and Hi-C assisted assembly of *Elaeagnus angustifolia* L., a deciduous tree belonging to *Elaeagnaceae*

**DOI:** 10.1101/665927

**Authors:** Yunfei Mao, Qin Hu, Manman Zhang, Lu Yang, Lulu Zhang, Yunyun Wang, Yijun Yin, Huiling Pang, Yeping Liu, Xiafei Su, Song Li, XinXing Cui, Fengwang Ma, Naibin Duan, Donglin Zhang, Yanli Hu, Zhiquan Mao, Xuesen Chen, Xiang Shen

## Abstract

*Elaeagnus angustifolia* L. is a deciduous tree of the *Elaeagnaceae* family. It is widely used in the study of abiotic stress tolerance in plants and for the improvement of desertification-affected land due to its characteristics of drought resistance, salt tolerance, cold resistance, wind resistance, and other environmental adaptation. Here, we report the complete genome sequencing using the Pacific Biosciences (PacBio) platform and Hi-C assisted assembly of *E. angustifolia*. A total of 44.27 Gb raw PacBio sequel reads were obtained after filtering out low-quality data, with an average length of 8.64 Kb. And 39.56 Gb clean reads was obtained, with a sequencing coverage of 75×, and Q30 ratio > 95.46%. The 510.71 Mb genomic sequence was mapped to the chromosome, accounting for 96.94% of the total length of the sequence, and the corresponding number of sequences was 269, accounting for 45.83% of the total number of sequences. The genome sequence study of *E. angustifolia* can be a valuable source for the comparative genome analysis of the *Elaeagnaceae* family members, and can help to understand the evolutionary response mechanisms of the *Elaeagnaceae* to drought, salt, cold and wind resistance, and thereby provide effective theoretical support for the improvement of desertification-affected land.

## Introduction

*Elaeagnus angustifolia* L., also known as silver willow and cinnamon, is a deciduous tree belonging to the *Elaeagnaceae* family (Fig. 1). It is native to central and western Africa and is distributed in the United States, Canada, the Mediterranean coast, southern Russia, Iran, and India. It shows a wide distribution area in China, where is is distributed in the Xinjiang, Gansu, Ningxia, Inner Mongolia, and other provinces(Wang *et al.*, 2014). The fruit, branches, leaves, and flowers of *E. angustifolia* can be used as medicine owing to multiple beneficial properties. The fruit is rich in sugars, flavonoids, and other substances that can regulate the blood circulation of the human body and improve the immunity of the body; the branches, leaves, and flowers are beneficial for anti-aging, and treatment of burns, bronchitis, dyspepsia, and neurasthenia(Min *et al.*,2006; Vitas *et al.*, 2004; Wang *et al.*,2006). The flowers are also used for extracting aromatic oil, which is used as a flavoring raw material in soap(Liu *et al.*, 2003).

**Figure 1.**
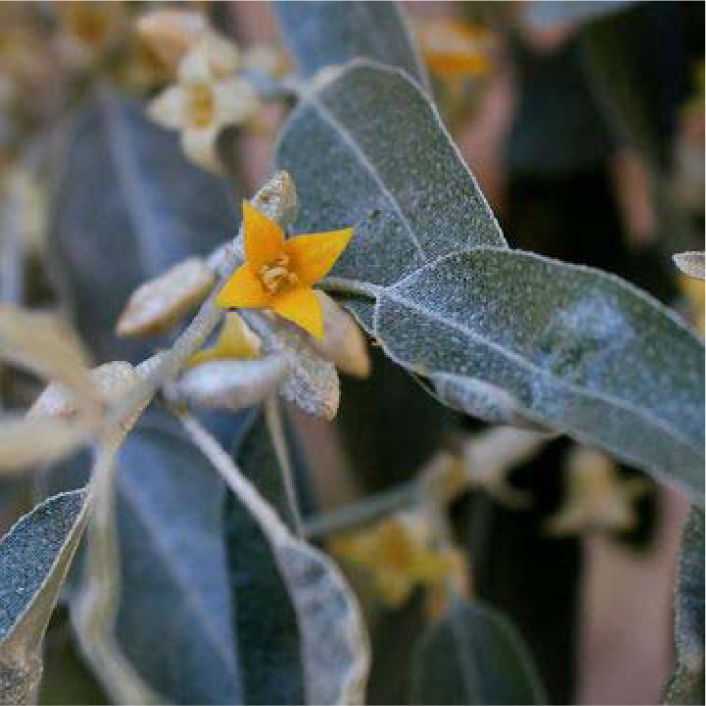
Elaeagnus angustifolia

At present, land desertification is a serious global phenomenon. Due to economic development needs, the effects of various methods such as terraced fields and grazing control to recover from land desertification are not significant in Spain, Greece, Turkey and other countries(Salvati *et al.*, 2016). *E. angustifolia* shows the characteristics of drought resistance, salt tolerance, cold resistance, wind resistance, easy reproduction, and strong adaptability(Huang *et al.*, 2005). The root rhizobium has important effects on nitrogen fixation and soil improvement, which can reform saline-alkali land and improve desertification-affected land(Liu, 2015). In recent years, *E. angustifolia* has been cultivated in Hebei, Heilongjiang, Henan, Shanxi, Shandong, and other provinces in China(Guo *et al.*, 2008).

Although the nrDNA ITS sequence data of *Elaeagnaceae* are abundant in the GenBank at present(He, 2012), studies on genome sequencing of *Elaeagnaceae* have not yet been reported, and the genome is an important basis for analyzing the evolution of *Elaeagnaceae*. At present, Pacific BioSciences(PacBio) technology, a third-generation sequencing technology, and Hi-C assisted assembly technology have become increasingly reliable and the genome sequencing has been completed for *Saccharum spontaneum* L.(Zhang *et al.*, 2018) and *Ammopiptanthus nanus*(Gao *et al.*, 2018).

In this study, we applied PacBio technology and Hi-C assisted assembly technology to sequence the genome of *E. angustifolia*, which is a valuable source for comparative genomic analysis of the *Elaeagnaceae* family members. Genome sequencing can help understand the response mechanism of the *Elaeagnaceae* to drought, salt, cold and wind resistance, and provide an effective theoretical basis for planting *E. angustifolia* to recover from global land desertification.

## Materials and Methods

### Sample collection

Samples from an *Elaeagnus angustifolia* L. tree (imported from Xinjiang province, NCBI Taxonomic ID, 36777) were collected from the south campus of Shandong Agricultural University for genomic DNA sequencing, and Hi-C assisted assembly.

### Genomic DNA sequencing and Hi-C assisted assembly

After collection, tissues were immediately immersed in liquid nitrogen and stored until DNA extraction. DNA was extracted using the Cetyltrimethyl Ammonium Bromide (CTAB) method. The quality of the extracted genomic DNA was checked using 1% agarose gel electrophoresis, and the concentration was quantified using a Qubit fluorimeter (Invitro-gen, Carlsbad, CA, USA). After checking the quantity and quality of the DNA sample, the library was constructed as shown in Supplementary Figure S1 in the order from left to right as shown in Supplementary Figure S2.

## Results and discussion

### Genomic results and statistics

We constructed two 270-bp libraries using genomic DNA of *E. angustifolia* samples. A total of 60.15 Gb of high-quality data was sequenced and filtered on Illumina Hiseq sequencing platform (San Diego, CA, USA), and the total sequencing depth was about 131×, which met the sequencing requirement of more than 50× (Supplementary Table S1). A total of 5,125,675 subreads were obtained by filtering low-quality data, and a total of 44.27 Gb raw PacBio sequel reads were obtained, with an average length of 8.64 kb (Supplementary Table S2). The subread N50 was 12,635 bp, and the average length was 8,636 bp (Supplementary Table S3). Subreads were corrected and assembled by Canu(Koren *et al.*, 2017), and the estimated genome size was found to be 781.09 Mb and Contig N50 was 486.92 Kb (Supplementary Table S4).

A kmer map of k = 19 was constructed using the two 270-bp library data (Supplementary Figure S3), which was used to evaluate genome size, repeat sequence ratio, and heterozygosity. The highest peak in the kmer distribution curve was found at the k-mer depth of 111. The sequences with kmer depth more than twice of the corresponding depth of the main peak, i.e. kmer sequences with a depth greater than 223, were repetitive sequences. The sequence with kmer depth appearing at half of the depth corresponding to the main peak, i.e. the kmer sequence with depth appearing around 55 was a heterozygous sequence. The total number of kmer obtained from sequencing data was 52,917,129,364. After removing those with depth abnormality, a total of 51,064,317,165 kmer sequences were used for the estimation of genome length, whose calculated length was about 456.24 Mbp. Based on distribution of kmer, the genome of this species was found to be a complex genome with high heterozygosity, with the content of repeat sequences estimated to be about 39.24%, and the degree of heterozygosity estimated to be about 1.47%.

Due to the relatively low conservation of repeat sequences among species, it is necessary to construct a specific repeat sequence database for the prediction of repeat sequences for specific species. With the help of LTR FINDER v1.05(Xu *et al.*, 2007), MITE Hunter(Han *et al.*, 2010), RepeatScout v1.0.5(Price *et al.*, 2005), and piler-df v2.4(Edgar *et al.*, 2005), the repeat sequence database of E. angustifolia genome was constructed based on the structure prediction and the principle of de novo prediction. The database was classified by PASTEClassifier(Wicker *et al*., 2007), and then merged with the database of Repbase(Jurka *et al*., 2005) as the final repetitive sequence database, and then repeated sequences were identified based on the constructed repeat sequence database using RepeatMasker v4.0.6(Tarailo-Graovac *et al*., 2009) software. The prediction yielded a repeat of about 263.44 Mb, accounting for 50.01%. The detailed prediction results are shown in Table 1.

**Table 1.**
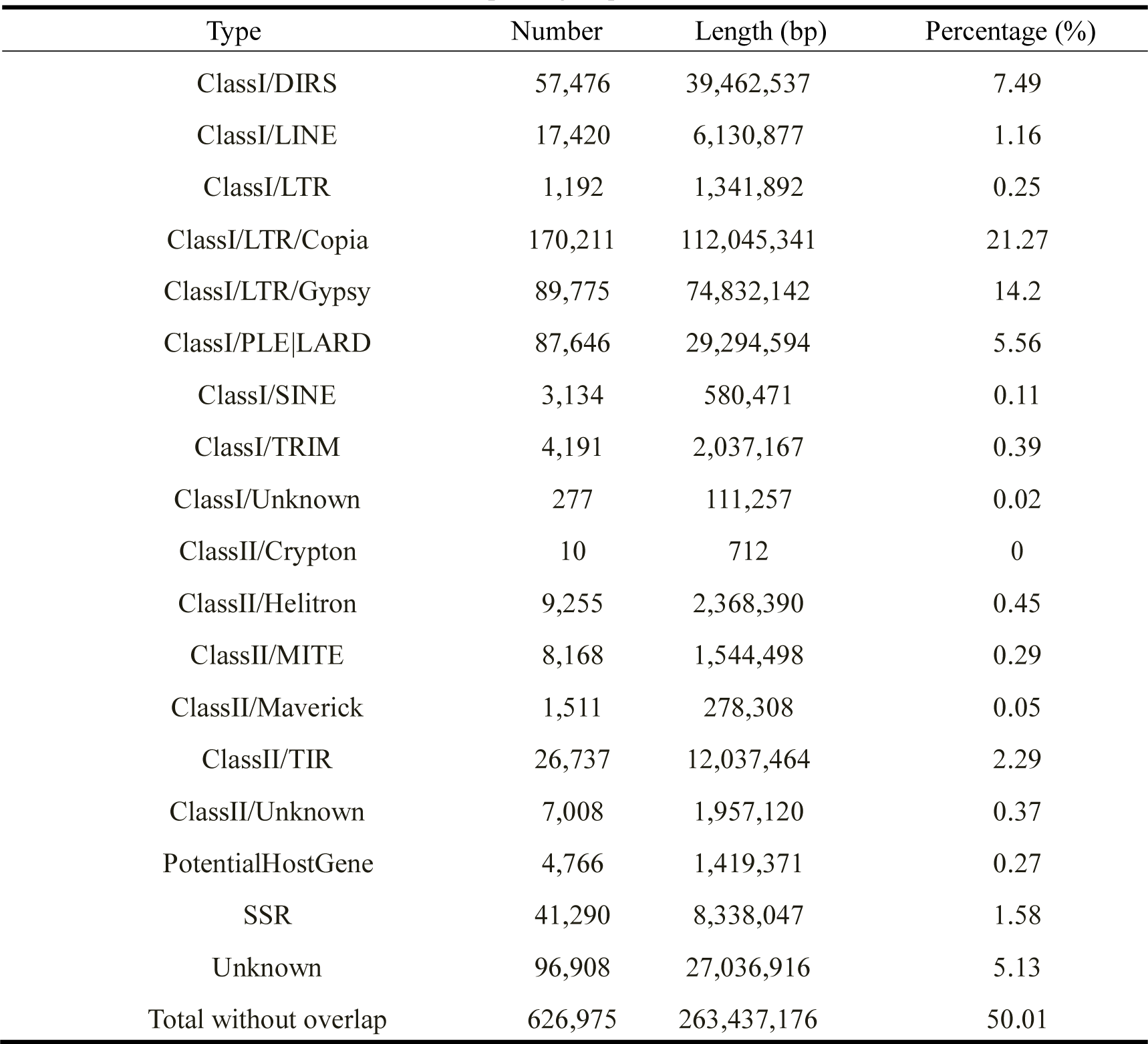
Repeating sequence statistics

| Type | Number | Length (bp) | Percentage (%) |
| --- | --- | --- | --- |
| ClassI/DIRS | 57,476 | 39,462,537 | 7.49 |
| ClassI/LINE | 17,420 | 6,130,877 | 1.16 |
| ClassI/LTR | 1,192 | 1,341,892 | 0.25 |
| ClassI/LTR/Copia | 170,211 | 112,045,341 | 21.27 |
| ClassI/LTR/Gypsy | 89,775 | 74,832,142 | 14.2 |
| ClassI/PLE/LARD | 87,646 | 29,294,594 | 5.56 |
| ClassI/SINE | 3,134 | 580,471 | 0.11 |
| ClassI/TRIM | 4,191 | 2,037,167 | 0.39 |
| ClassI/Unknown | 277 | 111,257 | 0.02 |
| ClassII/Crypton | 10 | 712 | 0 |
| ClassII/Helitron | 9,255 | 2,368,390 | 0.45 |
| ClassII/MITE | 8,168 | 1,544,498 | 0.29 |
| ClassII/Maverick | 1,511 | 278,308 | 0.05 |
| ClassII/TIR | 26,737 | 12,037,464 | 2.29 |
| ClassII/Unknown | 7,008 | 1,957,120 | 0.37 |
| PotentialHostGene | 4,766 | 1,419,371 | 0.27 |
| SSR | 41,290 | 8,338,047 | 1.58 |
| Unknown | 96,908 | 27,036,916 | 5.13 |
| Total without overlap | 626,975 | 263,437,176 | 50.01 |

TopHat(Trapnell *et al*., 2009) was used to compare the raw transcriptome data with the genome of E. angustifolia, and the number of bases in the Exon, Intron, and Intergenic regions were separately counted to evaluate the results of the gene prediction (Supplementary Table S5). The prediction of the genetic structure of E. angustifolia mainly used de novo prediction, homologous species prediction, and Unigene prediction, and then integrated the prediction results using EVM v1.1.1(Haas *et al*., 2008) software. Genscan(Burge *et al*., 1997), Augustus v2.4(Stanke *et al*., 2003), GlimmerHMM v3.0.4(Majoros *et al*., 2004), GeneID v1.4(Blanco *et al*., 2007), SNAP (version 2006-07-28) (Korf, 2004) were used for head-to-head prediction. GeMoMa v1.3.1(Keilwagen *et al*., 2016) was used for de novo prediction. His v2.0.4(Pertea *et al*., 2016) and Stringtie v1.2.3(Pertea *et al*., 2016) were used for assembly based on reference transcript, and TransDecoder v2.0(Haas *et al*., 2016)and gene marks-t v5.1(Tang *et al*., 2015) was used for gene prediction. PASA v2.0.2(Campbell *et al*., 2006) was used to predict the Unigene sequences without reference assembly based on transcriptome data. Finally, EVM v1.1.1(Haas *et al*., 2008) was used to integrate the prediction results obtained by the above three methods, and 31,730 genes were obtained after modification with PASA v2.0.2. The specific predicted information is shown in Table 2 and Supplementary Table S6. The number of genes supported by the three prediction methods was integrated, as shown in Supplementary Figure S4. As shown, the number of genes supported by homologous prediction and transcriptome prediction resulted in 30,771 genes, accounting for 96.98%, indicating the high prediction quality. At the same time, according to the gene function annotation, 96.89% of the genes could be annotated into NR and other databases, which further indicated that the gene prediction was reliable.

**Table 2.**
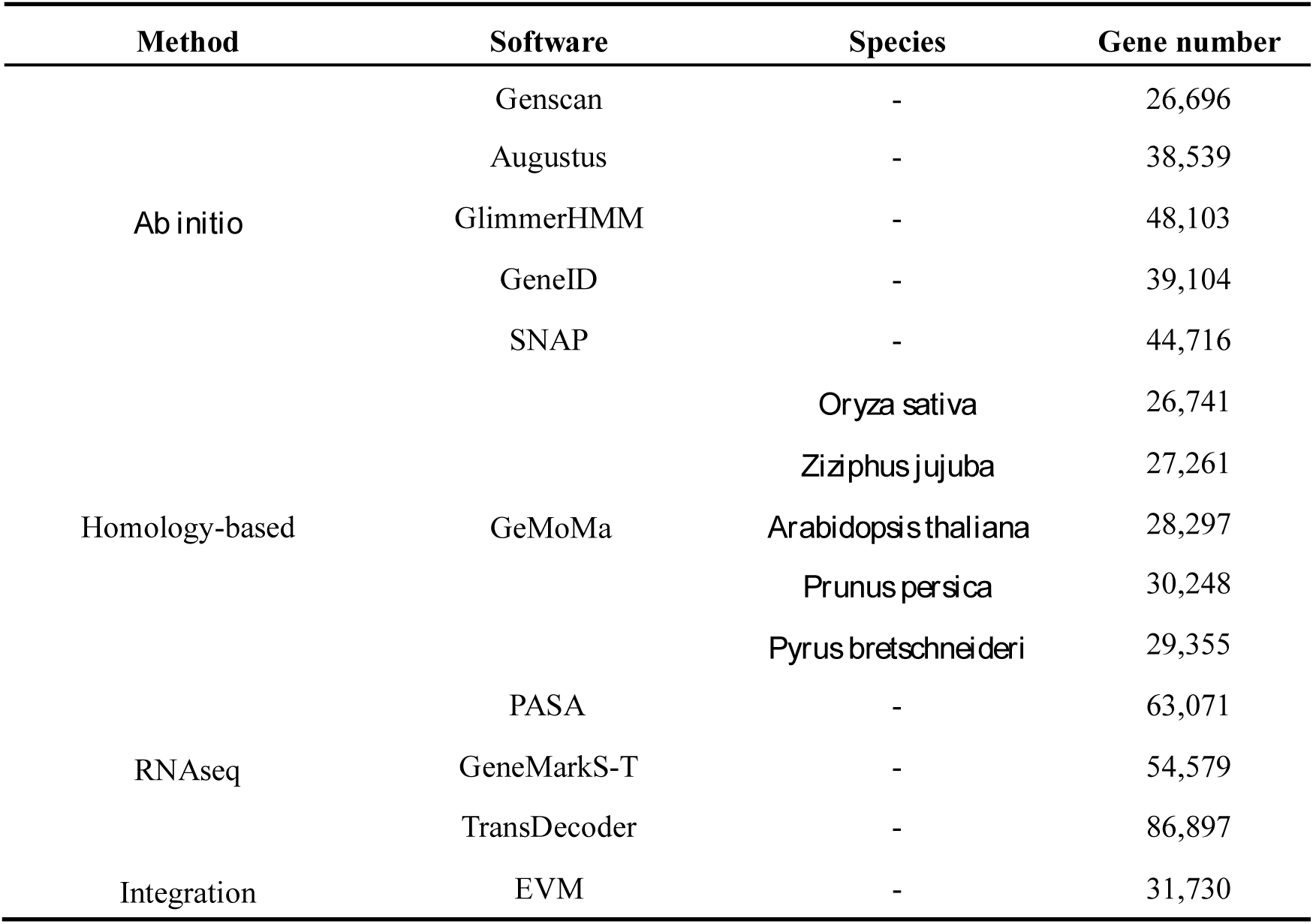
Gene prediction result statistics

BLAST v2.2.31(Birney *et al*., 2004) with an E-value cutoff of 1E-5 was used to align the predicted gene sequences with functional databases such as NR(Griffiths-Jones *et al*., 2005), KOG(Griffiths-Jones *et al*., 2006), GO(Nawrocki *et al*., 2013), KEGG(Lowe *et al*., 1997), and TrEMBL(She *et al*., 2009). Functional annotation analyses, namely the KEGG pathway annotation analysis, KOG functional annotation analysis, and GO functional annotation analysis were performed. A total of 30,743 of the predicted genes were annotated into databases such as the NR (Supplementary Table S7). By comparison with GenBlastA v1.0.4(She *et al*., 2009), homologous gene sequences were found in the genome with the true locus screened. GeneWise v2.4.1(Birney *et al*., 2004) was used to find immature termination codons and frame-shift mutations in the gene sequences, and pseudogenes were identified. A total of 2,173 pseudogenes were predicted (Supplementary Table S8).

### Hi-C assisted assembly

Based on Sequencing By Synthesis (SBS) technology, the Illumina high-throughput sequencing platform was used to sequence the Hi-C library to produce a large number of high-quality reads. Raw data for sequencing samples included two FASTQ files, including reads measured at both ends of all Hi-C constructed library fragments (Supplementary Figure S5). We obtained 39.56 Gb clean reads, with sequencing coverage of 75×, and Q30 ratio of > 95.46% (Supplementary Table S9).

BWA(Li *et al*., 2009) and SAMtools (version: 0.7.10-r789) were used to map the pair-end data with the assembled genome sequence. The ratio of reads mapped to the assembled genome was 90.68%, and the ratio of Unique Mapped Read Pairs was 61.13%, indicating that the Hi-C data were good enough for subsequent analysis (Supplementary Table S10). We used HiC-Pro(Servant *et al*., 2015) to filter and evaluate the Hi-C data. The Invalid Interaction Pairs ratio cannot exceed 80% if it is a usable Hi-C library(Belton *et al*., 2012). Invalid Interaction Pairs mainly include Self-circle Ligation, Dangling Ends type, Re-ligation type, and other discarded types(Belton *et al*., 2012; Hu *et al*., 2013; Imakaev *et al*., 2012; Lajoie *et al*., 2015; Servant *et al*., 2015). A total of 80.79 M pairs of reads on the genome were obtained in this experimental library. Among them, 72.97 M pairs were valid Hi-C data, accounting for 90.32% of the data on the genome, and the ratio of Invalid Interaction Pairs was 9.68% (Supplementary Table S11).

After Hi-C assembly, a total of 51.71 Mb of genomic sequence was mapped to the chromosome, accounting for 96.94% of the total length of the sequence, and the corresponding number of sequences was 269, accounting for 45.83% of the total number of sequences. Among the sequences located on the chromosome, the sequence length that could determine the order and direction was 473.91 Mb, accounting for 92.8% of the total length of the sequence located on the chromosome, and the number of corresponding sequences was 104, accounting for 38.66% of the total number of sequences located on the chromosome (Supplementary Table S12).

For Hi-C assembled into the genome of the chromosome, the length was cut into a bin of 100 Kb, and then the number of Hi-C Read Pairs was covered between any two bins as the intensity signal of the interaction between the two Bins (Fig 2). A total of 14 chromosome groups could be clearly distinguished; within each group, it could be seen that the intensity of the interaction at the diagonal position was higher than that of the non-diagonal position, indicating that the interaction strength between adjacent sequences (diagonal position) in the result of Hi-C chromosome assembly was high, while that between non-adjacent sequences (non-diagonal position) was weak, which was consistent with the principle of Hi-C assisted genome assembly and proved that the genome assembly had a good effect.

**Fig 2.**
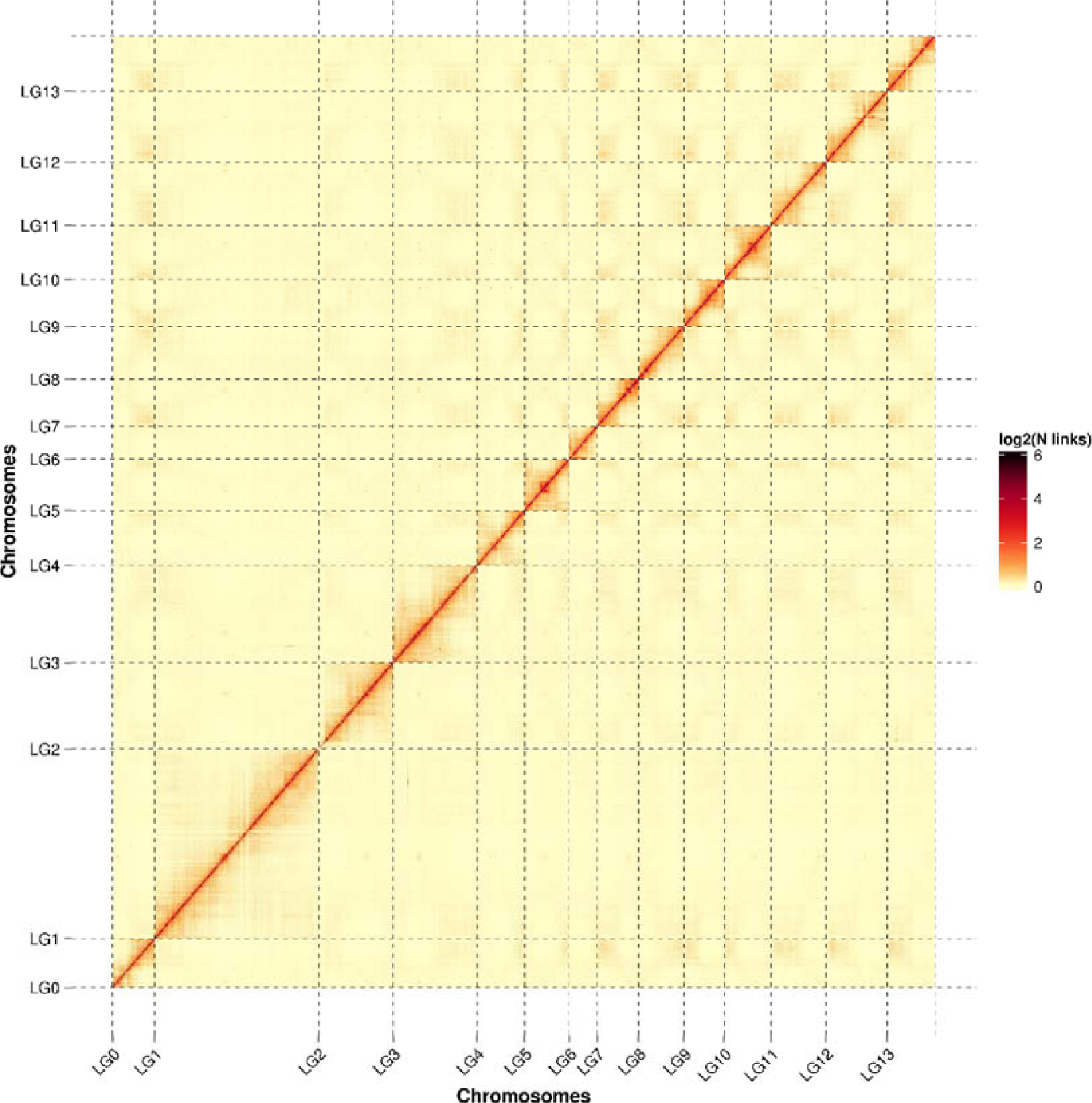
Hi-C assembly chromosome interaction heat map

## Conclusion

In this study, the genome of *Elaeagnus angustifolia* L. was obtained using PacBio technology, and Hi-C assisted assembly technology. Thus, our findings are a valuable source for comparative genomic analyses of the *Elaeagnaceae* and can help understand the response mechanism of the *Elaeagnaceae* to drought, salt, cold and wind resistance, thereby providing an effective theoretical basis for planting *E. angustifolia* to reverse global land desertification.

## Acknowledgements

We would like to thank Editage (www.editage.com) for providing linguistic assistance during the preparation of this manuscript. The research was supported by the National Science and Technology Support Program, China(NO1: 2014BAD16B02), the Shandong Key Research and Development Program, China(NO2: 2018GNC113019), the Fruit innovation team in Shandong Province, China(NO3: SDAIT-06-07), and the fund of National Modern Agro-industry Technology Research System of China(NO4: CARS-28).

## Author contributions

Yunfei Mao, Xinxing Cui, Yanli Hu and Xiang Shen planned and designed the research. Yunfei Mao, Qin Hu, Manman Zhang, Lu Yang, Lulu Zhang, Yunyun Wang, Yijun Yin, Huiling Pang, Yeping Liu, Xiafei Su and Song Li peformed experiments, conducted fieldwork, analysed data etc. Yunfei Mao, Fengwang Ma, Naibin Duan, Donglin Zhang, Yanli Hu, Zhiquan Mao, Xuesen Chen and Xiang Shen wrote the manuscript. Every author contributed equally.

## Supporting information

Additional Supporting information may be found in the online version of this article:

**Figure S1** DNA library components.

**Figure S2** Hi-C sequencing experiment process.

**Figure S3** Distribution of k-mers of length 19 from the Illumina Hiseq reads.

**Figure S4** The integrated gene is derived from the distribution map of three prediction methods.

**Figure S5** FASTQ file format.

**Table S1** Sample sequencing result statistics.

**Table S2** Length distribution of subreads of Pac-bio sequencing.

**Table S3** Filtering raw data of Pac-bio sequencing.

**Table S4** Genome assembly evaluation statistics.

**Table S5** Transcriptome comparison region statistics.

**Table S6** Gene information statistics.

**Table S7** Gene function annotation statistics.

**Table S8** Pseudogene Prediction Results.

**Table S9** Sequencing data volume statistics.

**Table S10** Clean data and genome alignment results statistics.

**Table S11** Hi-C sequencing data Validation.

**Table S12** Hi-C assembles data statistics.

## Reference

Belton, J.M., McCord, R.P., Gibcus, J.H., Naumova, N., Zhan, Y., Dekker, J. (2012) Hi-C: a comprehensive technique to capture the conformation of genomes. Methods. 58, 268–276.

Birney, E., Clamp, M., Durbin, R. (2004) GeneWise and genomewise. Genome research. 14, 988–995.

Birney, E., Clamp, M., Durbin, R. (2004) GeneWise and genomewise. Genome research. 14, 988–995.

Blanco, E., Parra, G., Guigó, R. (2007) Using geneid to identify genes. Current protocols in bioinformatics. https://doi.org/10.1002/0471250953.bi0403s18.

Burge, C., Karlin, S. (1997) Prediction of complete gene structures in human genomic DNA. Journal of molecular biology. 268, 78–94.

Campbell, M.A., Haas, B.J., Hamilton, J.P., Mount, S.M., Buell, C.R. (2006) Comprehensive analysis of alternative splicing in rice and comparative analyses with Arabidopsis. BMC genomics. 7, 327.

Edgar, R.C., Myers, E.W. (2005) PILER: identification and classification of genomic repeats. Bioinformatics. 21, i152–i158.

Gao, F., Wang, X., Li, X.M., Xu, M.Y., Li, H.Y., Abla, M., Sun, H.G., Wei, S.J., Feng, J.C., Zhou, Y.J. (2018) Long-read sequencing and de novo genome assembly of Ammopiptanthus nanus, a desert shrub. GigaScience. 7, 1–5.

Griffiths-Jones, S., Grocock, R.J., Van Dongen, S., Bateman, A., Enright, A.J. (2006) miRBase: microRNA sequences, targets and gene nomenclature. Nucleic acids research. 34, D140–D144.

Griffiths-Jones, S., Moxon, S., Marshall, M., Khanna, A., Eddy, S.R., Bateman, A. Rfam: annotating non-coding RNAs in complete genomes. Nucleic acids research. 33, D121–D124.

Guo, L.J., Wang, Y.T. (2008) Conservation Research and Prospects of Elaeagnus Germplasm Resources and Utilization Values. Chinese Wild Plant Resources. 27, 32–34.

Haas, B.J., Papanicolaou, A. (2016) TransDecoder (Find Coding Regions Within Transcripts). http://transdecoder.github.io.

Haas, B.J., Salzberg, S.L., Zhu, W., Pertea, M., Allen, J.E., Orvis, J., White, O., Buell, C.R., Wortman, J.R. (2008) Automated eukaryotic gene structure annotation using EVidenceModeler and the Program to Assemble Spliced Alignments. Genome Biol. 9, R7.

Han, Y., Wessler, S.R. (2010) MITE-Hunter: a program for discovering miniature inverted-repeat transposable elements from genomic sequences. Nucleic acids research. gkq862.

He, Y.H. (2012) Bioinformatic Analysis of the Elaeagnaceae nrDNA ITS Sequences. Northwest Normal University, China.

Hu, M., Deng, K., Qin, Z.H., Liu, J.S. (2013) Understanding spatial organizations of chromosomes via statistical analysis of Hi-C data. Quantitative Biology. 1, 156–174.

Huang, J.H., Maimaitijiang, Yang C.H., Wang C.F. (2005) Present Situation and Prospect about the Study of *Elaeagnus angustifolia* L. Chinese Wild Plant Resources. 24, 26–29, 33.

Imakaev, M., Fudenberg, G., McCord, R.P., Naumova, N., Goloborodko, A., Lajoie, B.R., Dekker, J., Mirny, L.A. (2012) Iterative correction of Hi-C data reveals hallmarks of chromosome organization. Nat Methods. 9, 999–1003.

Jurka, J., Kapitonov, V.V., Pavlicek, A., Klonowski, P., Kohany, O., Walichiewicz, J. (2005) Repbase Update, a database of eukaryotic repetitive elements. Cytogenetic and genome research. 110, 462–467.

Keilwagen, J., Wenk, M., Erickson, J.L., Schattat, M.H., Jan, G., Frank, H. (2016) Using intron position conservation for homology-based gene prediction. Nucleic acids research. 44, e89–e89.

Koren, S., Walenz, B.P., Berlin, K., Miller, J.R., Bergman, N.H., Phillippy, A.M. (2017) Canu: scalable and accurate long-read assembly via adaptivemer weighting and repeat separation. Genome Res. 27, 722–736.

Korf, I. (2004) Gene finding in novel genomes. BMC bioinformatics. 5, 59.

Lajoie, B.R., Dekker, J., Kaplan, N. (2015) The Hitchhiker’s guide to Hi-C analysis: practical guidelines. Methods. 72, 65–75.

Li, H., Durbin, R. (2009) Fast and accurate short read alignment with Burrows-Wheeler transform. Bioinformatics. 25, 1754–1760.

Lin, M., Todoric, D., Mallory, M., Luo B.S., Trottier, E., Dan, H.H. (2006) Monoclonal antibodies binding to the cell surface of *Listeria monocytogenes* serotype 4b. Journal of Medical Microbiology. 55, 291–299.

Liu, Y.W., Di, D.L., Wang, Q. (2003) Study on Chemical Components and Fingerprint of Volatile Oil from Flowers of *Elaeagnus angustifolia* L. Food Science. 24, 111–113.

Liu, Y.Z. (2015) Studies on Genetic Diversity of Elaegnus angustifolia L.. Hunan Agricultural University, China.

Lowe, T.M., Eddy, S.R. (1997) tRNAscan-SE: a program for improved detection of transfer RNA genes in genomic sequence. Nucleic acids research. 25, 0955–0964.

Majoros, W.H., Pertea, M., Salzberg, S.L. (2004) TigrScan and GlimmerHMM: two open source ab initio eukaryotic gene-finders. Bioinformatics. 20, 2878–2879.

Nawrocki, E.P., Eddy, S.R. (2013) Infernal 1.1: 100-fold faster RNA homology searches. Bioinformatics. 29, 2933–2935.

Pertea, M., Kim, D., Pertea, G.M., Leek, J.T., Salzberg, S.L. (2016) Transcript-level expression analysis of RNA-seq experiments with HISAT, StringTie and Ballgown. Nature Protocols. 11, 1650.

Price, A.L., Jones, N.C. (2005) Pevzner PA: De novo identification of repeat families in large genomes. Bioinformatics. 21, i351–i358.

Salvati, L., Kosmas, C., Kairis, O., Karavitis, C., Acikalin, S., Belgacem, A., Solé-Benet, A., Chaker, M., Fassouli, V., Gokceoglu, C., Gungor, H., Hessel, R., Khatteli, H., Kounalaki, A., Laouina, A., Ocakoglu, F., Ouessar, M., Ritsema, C., Sghaier, M., Sonmez, H., Taamallah, H., Tezcan, L., de Vente, J., Kelly, C., Colantoni, A., Carlucci, M. (2016) Assessing the effectiveness of sustainable land management policies for combating desertification: A data mining approach. Journal of Environmental Management. 183, 754–762.

Servant, N., Varoquaux, N., Lajoie, B.R., Viara, E., Chen, C.J., Vert, J.P., Heard, E., Dekker, J., Barillot, E. (2015) HiC-Pro: an optimized and flexible pipeline for Hi-C data processing. Genome Biology. 16, 1–11.

She, R., Chu, J.S.C., Wang, K., Pei, J., Chen, N.S. (2009) GenBlastA: enabling BLAST to identify homologous gene sequences. Genome Research. 19, 143.

Stanke, M., Waack, S. (2003) Gene prediction with a hidden Markov model and a new intron submodel. Bioinformatics. 19, ii215–ii225.

Tang, S., Lomsadze, A., Borodovsky, M. (2015) Identification of protein coding regions in RNA transcripts. Nucleic Acids Research. 43, e78.

Tarailo-Graovac, M., Chen, N. (2009) Using RepeatMasker to identify repetitive elements in genomic sequences. Current Protocols in Bioinformatics. https://doi.org/10.1002/0471250953.bi0410s25.

Trapnell, C., Pachter, L., Salzberg, S.L. (2009) TopHat: discovering splice junctions with RNA-Seq. Bioinformatics. 25, 1105–1111.

Vitas, A.I., eAguado, V. I.G.-J. (2004) Occurrence of *Listeria monocytogenes* in fresh and processed foods in Navarra (Spain). International Journal of Food Microbiology. 90, 349 –356.

Wang, B.S., Qu, H.Y., Ma, J., Sun, X.L., Wang D., Zheng Q.S., Xing D. (2014) Protective effects of elaeagnus angustifolia leaf extract against myocardial ischemia/reperfusion injury in isolated rat heart. Journal of Chemistry. 2014, 1–6.

Wang, Y., Zhao, P., Wang, Y.L., Zhang Y. (2006) Nutritional composition of wild Elaeagnus angustifolia fruits. Journal of Gansu Agricultural University. 41, 130–132.

Wicker, T., Sabot, F., Hua-Van, A., Bennetzen, J.L., Capy, P., Chalhoub, B., Flavell, A., Leroy, P., Morgante, M., Panaud, O. (2007) A unified classification system for eukaryotic transposable elements. Nature Reviews Genetics. 8, 973–982.

Xu, Z., Wang, H. (2007) LTR_FINDER: an efficient tool for the prediction of full-length LTR retrotransposons. Nucleic Acids Research. 35, W265–W268.

Zhang, J.S., Zhang X.T., Tang, H.B., Zhang Q., Hua, X.T., Ma, X.K., Zhu, F., Jones, T. X.G., Zhu, Bowers J., Wai, C.M., Zheng, C.F., Shi, Y., Chen, S., Xu, X.M., Yue, J.J., Nelson, D.R., Huang, L.X., Li, Z., Xu, H.M., Zhou, D., Wang, Y.J., Hu, W.C., Lin, J.S., Deng, Y.J., Pandey, N., Mancini, M., Zerpa, D., Nguyen, J.K., Wang, L.M., Yu, L., Xin, Y.H., Ge, L.F., Arro, J., Han, J.O., Chakrabarty, S., Pushko, M., Zhang, W.P., Ma, Y.H., Ma, P.P., Lv, M.J., Chen, F.M., Zheng, G.Y. Xu, J.S., Yang, Z.H., Deng, F., Chen, X.Q., Liao, Z.Y., Zhang, XX, Lin, Z.C., Lin, H., Yan, H.S., Kuang, Z., Zhong, W.M., Liang, P.P., Wang, G.F., Yuan, Y., Shi, J.X., Hou, J.X., Lin, J.X., Jin, J.J., Cao□, P.J., Shen, Q.C., Jiang, Q., Zhou, P., Ma, YY, Zhang, X.D., Xu, R.R., Liu, J., Zhou, Y.M., Jia, H.F., Ma, Q., Qi, R., Zhang, Z.L., Fang, J.P., Fang, H.K., Song, J.J., Wang, M.J., Dong, G.G., Wang, G., Chen, Z., Ma, T., Liu, H., Dhungana, S.R., Huss, S.E., Yang, X.P., Sharma, A., Trujillo, J.H., Martinez, M.C., Hudson, M., Riascos, J.J., Schuler, M., Chen, L.Q., Braun, D.M., Li, L., Yu, Q.Y., Wang□, J.P., Wang, K., Schatz, M.C., Heckerman, D., Sluys, M.-A.V., Souza, G.M., Moore, P. H., Sankoff, D., Buren, R.V., Paterson, A.H., Nagai, C., Mingll, R. (2018) Allele-defined genome of the autopolyploid sugarcane *Saccharum spontaneum* L.. Nature Genetics. 50, 1565–1573.

